# Hidden variation in polyploid wheat drives local adaptation

**DOI:** 10.1101/217828

**Authors:** Laura-Jayne Gardiner, Ryan Joynson, Jimmy Omony, Rachel Rusholme-Pilcher, Lisa Olohan, Daniel Lang, Caihong Bai, Malcolm Hawkesford, David Salt, Manuel Spannagl, F.X.Klaus Mayer, John Kenny, Michael Bevan, Neil Hall, Anthony Hall

## Abstract

Wheat has been domesticated into a large number of agricultural environments and has a remarkable ability to adapt to diverse environments. To understand this process, we survey genotype, repeat content and DNA methylation across a bread wheat landrace collection representing global genetic diversity. We identify independent variation in methylation, genotype and transposon copy number. We show that these, so far unexploited, sources of variation have had a massive impact on the wheat genome and that ancestral methylation states become preferentially ‘hard coded’ as SNPs via 5-methylcytosine deamination. These mechanisms also drive local adaption, impacting important traits such as heading date and salt tolerance. Methylation and transposon diversity could therefore be used alongside single nucleotide polymorphism (SNP) based markers for breeding.

## Background

One of the most important questions in plant breeding is the nature of the genomic variation that has been selected for improving phenotypes. Although it is likely that all forms of genomic change contribute to performance variation and to hybrid vigour, the role of epigenetic variation in crop improvement is not well understood, despite being widespread and highly variable^1^. It is now clear that epigenetic variation can be stably inherited^2,3^ and that spontaneous epialleles are rare. Therefore, epigenetic variants could potentially be used in breeding programmes and their contributions to trait variation assessed alongside classical genetic variation. To identify new sources of variation for crop improvement, and to understand the contributions of variation to traits, it is important to assess both genomic and epigenetic variation in crop species.

Epigenetic states of genes in crop plants have been shown to have a major influence on traits. Gene body methylation (gbM) can influence splice-site efficiency by differential CHG methylation of splice acceptor sites, indicating that epiallelic variation can contribute to differential mRNA accumulation^4^. In domesticated polyploid cotton and wild relatives there is extensive epigenetic variation, with methylation differences between homoeologous genes. One example is *COL2D* that is repressed by methylation in wild relatives but is activated by loss of methylation in allotetraploid cotton, influencing flowering time in domesticated lines^5^. The causal gene of a major QTL enhancing resistance to maize stalk rot, *ZmCCT*, is in two epigenetic states. One has CACTA-like TE upstream of *ZmCCT* promoter and one without that has enriched methylated CG that supressed expression and increased disease susceptibility^6^. Similar mechanisms of epigenetic change in gene expression mediated by retrotransposons adjacent to promoters have also been noted in wheat^7^. Tissue-culture induced reduction in methylation of a retrotransposon in the intron of an oil palm *Deficiens* gene alters splicing and causes premature termination^8^. This epigenetic mechanism contributes to the mantled phenotype that limits clonal propagation of this key global crop.

Analyses of DNA methylation patterns in numerous plant accessions and species are starting to reveal the extent of epigenetic variation and the mechanisms involved in generating and maintaining it. Two general patterns of DNA methylation have been identified in plants, transposable element methylation patterns (teM) and gene-body methylation patterns (gbM). In *Arabidopsis thaliana* accessions it was shown that increased gbM is related to constitutive gene expression patterns, and that teM epialleles of genes tend to be expressed at lower levels. Geographic origin was a major predictor of DNA methylation levels and of altered gene expression caused by epialleles^9,10^. It is clear that natural epigenetic variation provides a source of phenotypic diversity alongside genetic variation however, currently, little is known about this epigenetic variation and its interaction with genetic diversity in hexaploid wheat populations.

The genomes of crop plants such as maize and wheat are mainly composed of massive tracts of diverse retroelements and DNA repeats that comprise up to 80% of the genome. These repeats are highly methylated to suppress expression and transposition to maintain genome stability^11^. Wheat is an allopolyploid, comprised of three independently maintained A, B and D sub-genomes that are functionally diploid^12^. Epigenetic mechanisms have been invoked to explain the emergence of key agronomic traits upon formation of hexaploid bread wheat, and to explain alterations in gene expression of homoeologous genes upon polyploidization^13^. Previously we showed that methylation patterns differ across the A, B and D sub-genomes and in broad terms reflected patterns of methylation of progenitor species^14^. Here we extend our analyses to a core collection of diverse bread wheat landraces in the Watkins collection^15^. Landraces are locally adapted wheat varieties that have not been subject to selective breeding, and represent a pool of diversity reflecting their wide adaptation to different growing environments. Such diversity is beginning to be used in breeding programmes, therefore it is timely to assess and understand both the genomic and epigenomic diversity in this population.

We identified three main sources of variation across wheat landraces; high transposable element (TE) variability, alongside epigenetic and genetic diversity. Although we found a general correlation between methylation patterns and genotypic variation, there was a geographical component to methylation patterns that may indicate a response to or selection by local environmental conditions. We also show that ancestral methylation states may become preferentially ‘hard coded’ as SNPs via 5-methylcytosine deamination. Finally, we show that tri-genome methylation is the most stable form of methylation, and genome specific methylation patterns correlate with gene expression differences between homoeologous genes.

## Results

### Methylation and genotype analysis across a wheat landrace diversity panel

To study epigenetic variation across gene-rich regions of the 17 Gb allohexaploid wheat genome, we used genomic enrichment (Agilent SureSelect) followed by bisulfite treatment and Illumina HiSeq paired-end sequencing. Capture probes were designed (12 Mb capture targeting 36 Mb) as described in our previous work^16^ (Supplementary Figure 1 from Olohan *et al,* 2017).

To accurately apply methyl-seq to a diversity panel we require bisulfite treated and untreated sequence data for each wheat accession to identify C-T SNPs, which would otherwise be incorrectly classified as unmethylated cytosines. This was achieved using a modified sequence capture protocol^16,17^ that generates two libraries for sequencing from one capture; a bisulfite-treated and an untreated library for each sample. Post-sequencing, the untreated datasets were aligned to the TGAC v1 Chinese Spring reference sequence and SNP calling was performed (Methods)^18^. We identified 716,018 SNPs on average per sample at ≥5X, of which, 316,767 were homozygous. Homozygous SNPs were used to correct the reference genome for each accession, this corrected reference was implemented for mapping the corresponding bisulfite-treated dataset (Methods).

Bisulfite-treated DNA from single seedlings was examined for 104 core lines from the Watkins landrace collection plus the reference variety Chinese Spring (Supplementary Table 1 and note 1a). We scored methylation at an average of 10.9M cytosines per sample (Supplementary Table 2 and note 1b) and across all samples, on average 98.7% of cytosine bases were successfully bisulfite converted (Supplementary Table 3 and note 1c).

### Genetic variation across the Watkins collection clusters geographically

From the 716,018 SNPs that were identified on average per sample, 53,341 SNP sites were identified across the 105 samples where; all samples showed mapping coverage at ≥5X and ≥1 sample had a SNP. For each SNP, the alternate allele frequency per sample was used for hierarchical clustering of the accessions (Methods). Using genotype information for sample clustering, accessions originating from Europe and the Mediterranean tend to cluster together while samples from larger geographic regions in Asia and Russia show higher diversity (Figure 1a and 1b).

**Figure 1.**
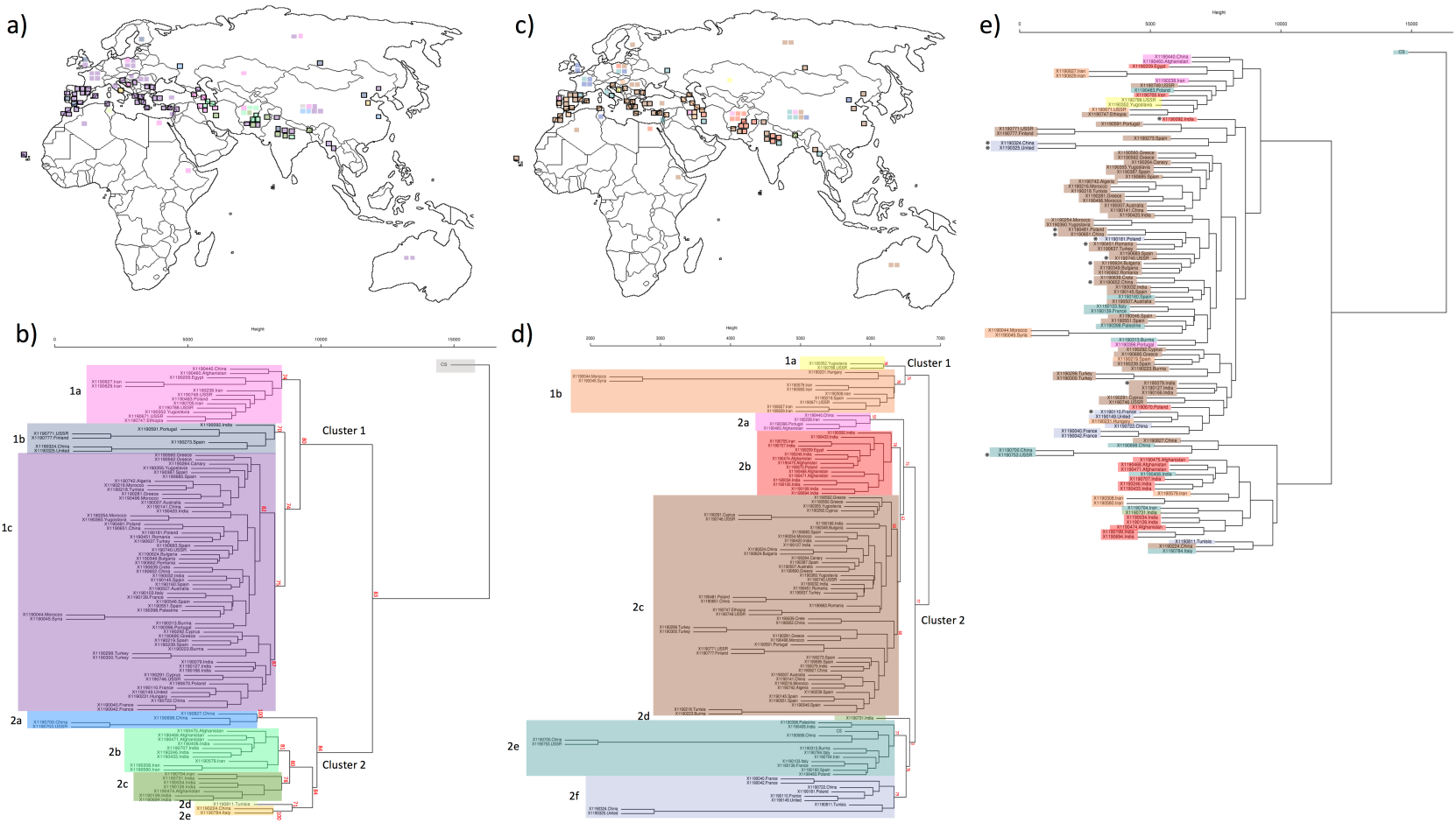
Geographical sample origins combined with hierarchical cluster analysis on 104 samples from the Watkins core collection plus Chinese Spring wheat. (**a**) Geographical positions of the samples colour coded by their allocated cluster from (**b**) after SNP hierarchical clustering. (**b**) Dendrogram constructed using the complete linkage method within the R package hclust to cluster samples based on SNP allele frequency across 53,341 SNP sites. The tree was cut into 8 groups (excluding the reference Chinese Spring) using the R package cutree and these clusters are colour-coded (Methods). (**c**) Geographical positions of the samples colour coded by their allocated cluster from (**d**) after CpG SMP hierarchical clustering. (**d**) Dendrogram constructed using the complete linkage method within the R package hclust to cluster samples based on methylation levels across 18,965 CpG SMP sites (taken from the 359,500 SMPs that were identified within the sample set). The tree was cut into 8 groups using the R package cutree and these clusters are colour-coded (Methods). (**e**) SNP based-dendrogram from (**b**) with individual samples colour-coded as per their cluster from the SMP-based dendrogram from (**d**). For geographical sample positions in (**a**) and (**c**) squares outlined in black represent samples with detailed positional information that is used for plotting, squares with no outline represent samples with only a country of origin. AU p-values were computed for the main clusters in (**b**) and (**d**) using the R package pvclust and are shown in red (Methods).

The Watkins collection clusters into two main ancestral groups; cluster 1 with 80 accessions (73.8% derived from Europe, Middle Eastern and South Mediterranean/African regions) while cluster 2 has 24 samples (87.5% mainly Asian) (Supplementary Table 4). This genotype-based population structure resembles that from previous analyses of the Watkins collection using array SNP data^15,19^ (see Supplementary note 2).

### SMPs are variable and cluster geographically across the Watkins collection

Global methylation patterns in Chinese Spring align closely to those of other plant species and previous analyses of Chinese Spring^14,20^ (Supplementary note 3; Supplementary Figure 1 and Supplementary Table 5). To assess epigenetic variation across the Watkins collection we identified 853,932 cytosines that were mapped to ≥10X in all 104 samples plus Chinese Spring. 359,500 (42.1%) of these cytosines were classified as single methylation polymorphism sites (SMPs) between the samples (Supplementary Table 6, Methods). Although methylation variability is high, the SMPs do not preferentially target any of the methylation contexts (CpG, CHG or CHH) (Supplementary Table 6).

0.5% of the 359,500 SMP sites show high methylation conservation between samples (methylated in ≥90%); these were mainly at CpG sites (86.2%) with a bias for transcribed regions (80.2%). Focusing on CpG sites, 13.9% of SMPs were methylated in ≥90 samples highlighting the increased stability of CpG sites compared to non-CpG sites. However, most SMPs (91.5%) are rare variants in <10% of the samples. Unlike highly conserved SMPs, these low-frequency SMPs show less bias for transcribed regions (74.2%) and increased bias for non-CpG sites potentially due to the more dynamic tissue specificity of this methylation (82.5% at CHH sites and 16.4% at CHG sites). Sample-specific SMPs were identified from the 359,500 SMPs (Methods); on average, each sample showed methylation at 26,980 SMP positions with a range of 11,279 to 64,659 SMPs per sample (Supplementary Table 7).

To analyze inter-sample variation in SMPs, for all 359,500 SMP sites, epi-allele frequency per sample was used for hierarchical clustering of accessions for CpG and non-CpG sites individually (Figure 2; Supplementary note 4; Supplementary Figure 2). When we order SMP sites by their total methylation across the accessions (vertical axes, Figure 2); for CpG sites there is a tendency for sites to show extremes of either high or low-level methylation, with typically more methylation in transcribed regions and less methylation in non-transcribed regions. Conversely, non-CpG SMP sites tend to show higher methylation in non-transcribed regions. Clustering the datasets by accession (horizontal axes, Figure 2); inter-sample variation is less obvious for non-CpG sites where most of the methylation is low-level or potentially tissue-specific (Figure 2c). However, more inter-sample methylation variation can be observed at CpG sites with both high and low-level methylation, therefore, accessions can be informatively compared (Figure 2a).

**Figure 2.**
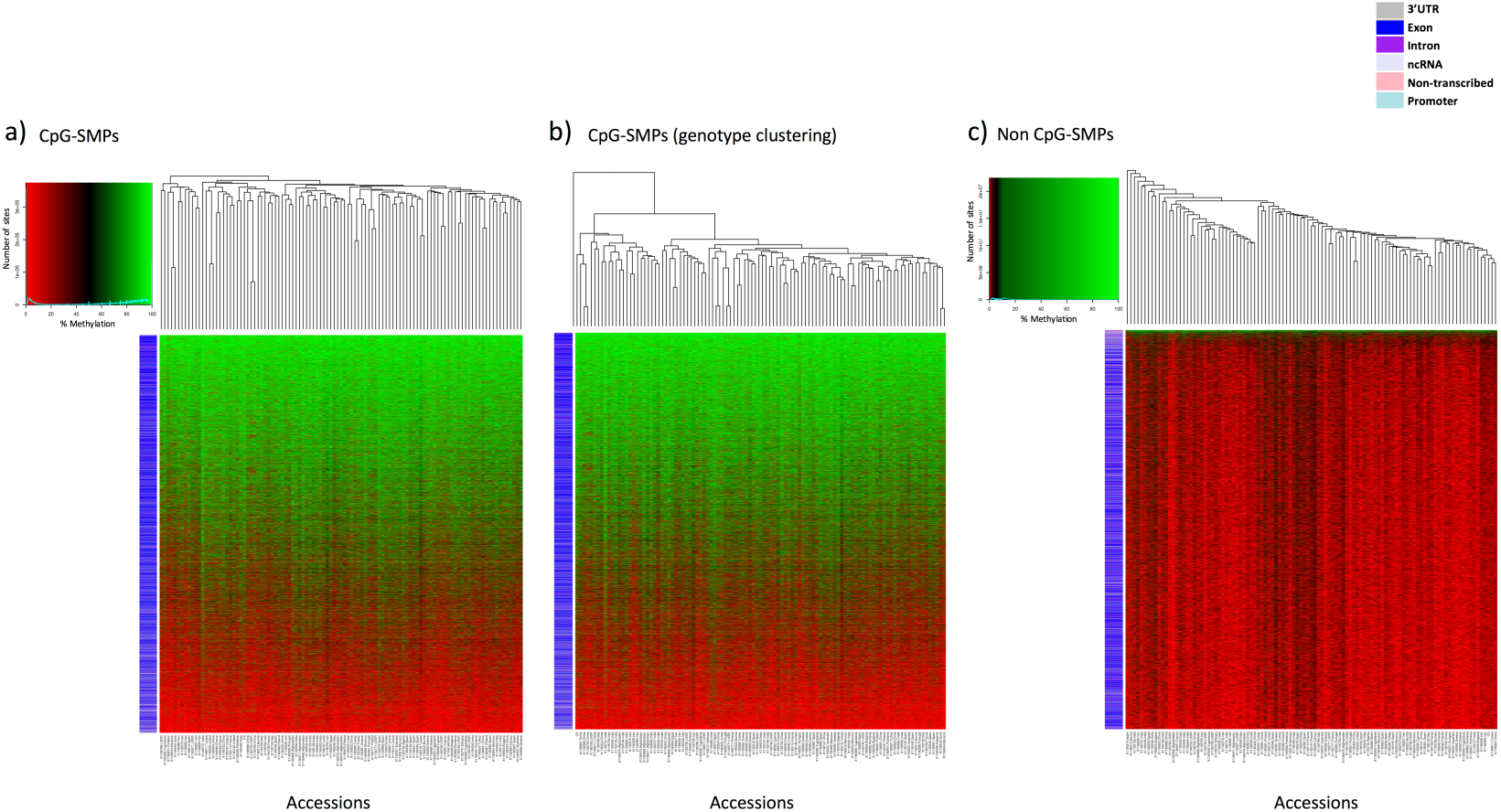
Visualizing methylation levels for the 105 wheat samples across 359,500 SMP sites. Using sites with coverage in all 104 Watkins collection accessions plus Chinese Spring we generated heatmaps for methylation levels across (**a**) CpG-SMPs (**b**) CpG-SMPs with accessions ordered by genotype using the heatmap from (**a**) with accessions re-ordered based on figure 1b’s SNP clustering dendrogram (shown on top horizontal axis) and (**c**) Non-CpG SMPs. Rows correspond to individual SMP sites and columns indicate accessions. The coloured row labels (barcodes) on the left of the heatmap indicate which genomic location a SMP falls into (see legend). SMP sites are ordered by their total methylation across the accessions on the vertical axes and accessions are clustered by SMP profiles on the horizontal axis (Methods).

Sample linkage across CpG methylation correlates with geographical sample proximity. Accessions from within the same country of origin tend to show higher linkage and cluster together closely, with 90.3% of the 31 regions analyzed containing a majority of samples (≥50%) from one linkage cluster (Figure 1c and 1d). From the top hierarchical level epigenetic population structure of the Watkins collection, samples cluster into two groups composed largely of accessions from mixed geographic locations; cluster 1 containing 12 samples derived from 50% Asian and 50% European/Middle Eastern locations and cluster 2 containing 93 samples derived from 41% Asian and 59% European/Middle Eastern locations (Figure 1; Supplementary Table 8). This population structure differs from the genotype-based population structure (Figure 1e); although both split into two sub-populations at the top hierarchical level, for genotype, these sub-populations showed one population from mixed geographic locations while the other was of Asian origin. We statistically compared the two cluster configurations (Figure 1b and 1d); the cluster configuration in the combined SMP and SNP trees (Figure 1e) was non-random (one-sample runs test with 39 runs: Z=-2.53, p = 0.011). This supports the existence of an association between the clustering patterns of SMPs and SNPs.

To determine the similarity of the epigenetic/genotypic profiles, frequency estimates were calculated for SNPs and SMPs across the genome. No correlation between genotype and epi-genotype was detected at this resolution (Supplementary Figure 3). We constructed distance matrices for the 18,965 CpG SMP sites and a comparably sized subset of the 53,341 variable SNP sites. Comparisons were then made using the non-parametric Mantel test to compute Pearson product-moment correlation between the matrices (Methods). A weak positive correlation of 0.394 was observed between the matrices (alpha=0.05, p<0.001) (Supplementary Figure 4). Since this correlation is low, genotype and methylation are likely to be linked but methylation can also develop independently of genetic variation. To corroborate this, we noted a broad-range tendency for samples clustering closely by SMP profile to show similar levels of methylation overall (Figure 2a; Supplementary Figure 5a). However, by ordering samples based on genotypic information and comparing their methylation profiles, only closely related samples share similar methylation levels (Figure 2b; Supplementary Figure 5b).

In summary, the methylation profiles of native accessions for mid/smaller sized countries, e.g. the UK, Greece, Afghanistan, Cyprus and Italy, are more likely to cluster together. These lines most likely evolved in similar environmental conditions and have similarly adapted methylation profiles. Conversely, we see samples from geographically distant locations with comparable methylation, this may represent conserved environmental conditions that have resulted in a similar adaptive change in methylation profiles. For samples where we have more accurate positional information for geographical origin, this association between methylation and local adaptation is clearer (see Supplementary Table 9; Supplementary note 5).

### Distinctive patterns of methylation are associated with different classes of gene function

Our analysis of the landraces clustered accessions with similar patterns of methylation into 8 distinct groups (Figure 1d). To assess if these clusters represented any functional consequences of gene methylation, genes that were methylated within each cluster were analysed by GO enrichment for molecular functions (topGO, p < 0.05). At this level of analysis, all 8 clusters had distinctive profiles of enriched GO terms across multiple functional categories of genes (Supplementary Table 10 and 11). To ascertain if there were any functional consequences of gene methylation patterns within these clusters, information on differential gene expression were included in these analyses, and are shown later in Supplementary Tables 24-26.

### Tri-genome is the most stable form of methylation

We classified methylation as tri-genome (in three sub-genomes), bi-genome and uni-genome (in two or one sub-genome respectively) (Methods, Supplementary Table 12). Supplementary Table 13 details differentially and tri-genome methylated CpG, CHH or CHG sites averaged across the samples. The observed methylation landscape largely reflects that seen in our previous analysis (Supplementary note 6)^14^.

To assess the relative stability of uni-, bi- and tri-genome methylation across the Watkins collection we identified positions that were uni-, bi- or tri-genome methylated in one or more of the samples. From these positions, we selected all sites that had mapping coverage ≥10X in all accessions, independent of their methylation status. Figure 3a highlights a median of 20.95% of accessions showing conserved tri-genome methylation compared to only 2.85% of accessions with conserved uni- or bi-genome methylation. Furthermore, 14.3% of tri-genome sites were methylated in the majority of samples (>= 90%) whereas, on average, only 1.08% of uni- and bi-genome sites showed methylation conservation on this scale (Supplementary Table 14). Tri-genome methylation is significantly more conserved across the accessions compared to uni and bi-genome methylation respectively (bi genome t=74.66, df=16508, p-value <2.2e-16; uni-genome t=67.56, df=17848, p-value <2.2e-16) and this tri-genome methylation is evenly distributed across the genome (Methods, Supplementary Figure 6; track 1, 5 and 9). Gene Ontology enrichments, for genes associated with the most stable subset of trigenome methylation (in >= 90% of samples), included core biological activities within the plant such as phosphorylation, intracellular transport, transcription regulation, oxidation-reduction, proteolysis and methylation (Supplementary Table 15).

**Figure 3.**
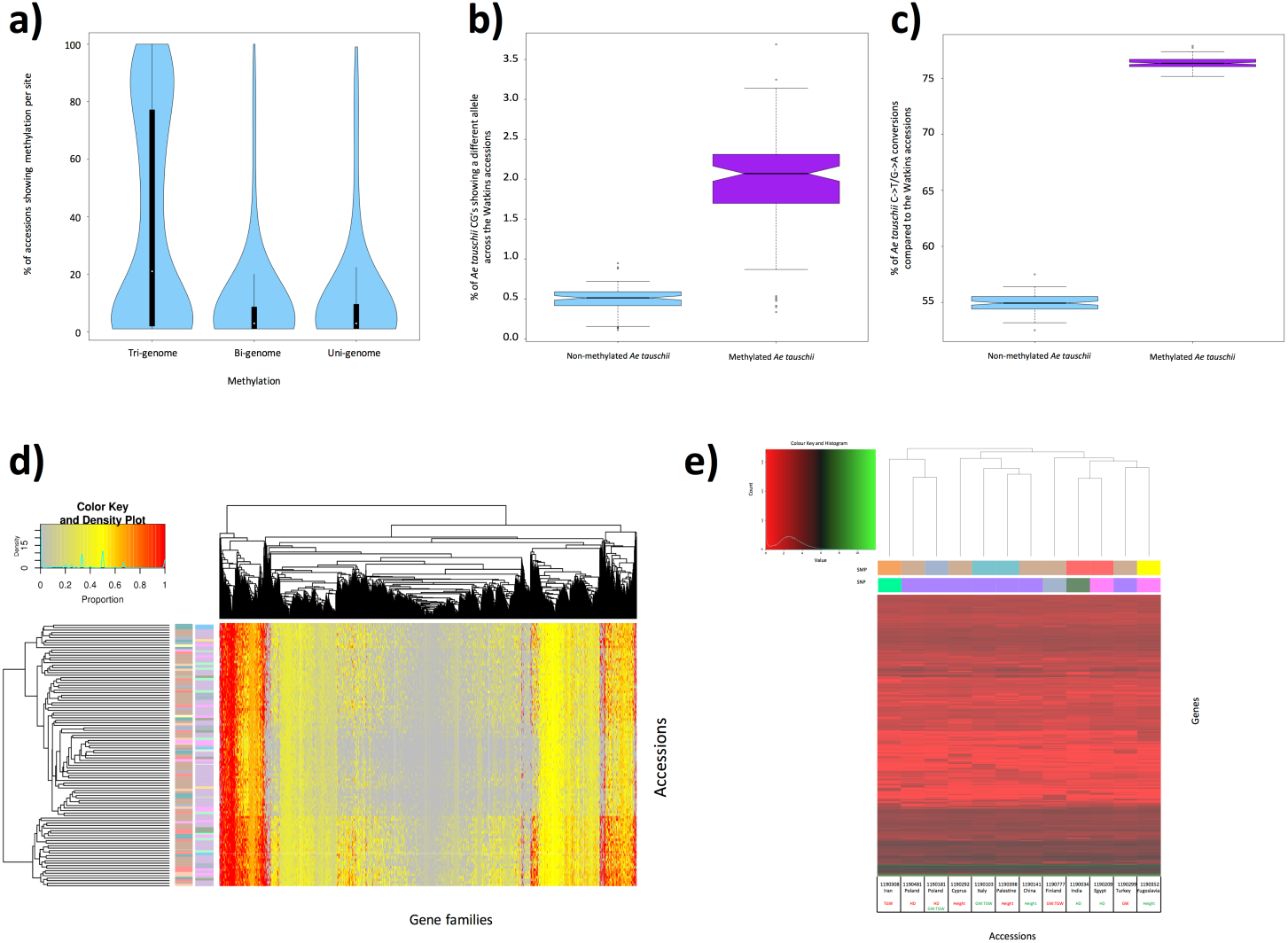
Analyzing methylation profiles across the Watkins collection. (**a**) Violin plots show the percentage of accessions showing methylation per analyzed site. Analyzed sites include Tri-genome, Bi-genome and Uni-genome methylated sites. A comparative subset of 11,769 sites was used for each category. (**b**) Ancestral methylation associates with an increased SNP rate. The percentage of methylated versus non-methylated *Ae. tauschii* cytosines that show a different allele in the Watkins. (**c**) Ancestral methylation demonstrates that 5-methylcytosines are preferentially deaminated to thymine. The percentage of methylated versus non-methylated *Ae. tauschii* cytosines with a C-to-T/G-to-A transition across the Watkins collection. (**d**) Sample clustering based on the gene families targeted by methylation. Many accessions from the same geographical origin show the same gene families targeted by methylation; thus, clustered close to each other in the Accessions axis (vertical dendrogram). Alongside the vertical dendrogram the two columns of row barcodes (left and right) correspond to the SMP clusters in Figure 1d and SNP clusters in Figure 1b respectively. (**e**) Sample clustering of the 12 accessions subjected to RNA-seq using average gene expression across the replicates for genes showing differential expression between at least 2 lines (after log2 transformation). Below the horizontal dendrogram the two barcode rows (top and bottom) correspond to the SMP and SNP clusters in Figure 1d and 1b respectively. Accessions are labelled by line number, country of origin and phenotype i.e. TGW (thousand grain weight), HD (heading date), GW (grain width) or Height with maximum values in green and minimum values in red.

### Genome-specific methylation associates with homoeologous SNPs

We analyzed methylation variation where all samples contained the same sequence. Looking at the cytosine residue sites that were mapped to ≥10X in all of the samples, most (89.0%) shared the same genetic sequence, i.e. cytosine CpG/CHG/CHH context, and were, therefore used to identify 359,500 SMPs. Methylation is a source of variation in the absence of genetic variation; however, we also assessed the impact of SNPs on methylation. Across all the samples, at cytosine sites showing tri-genome methylation, the average percentage of sites where a SNP altered the cytosine context between the sub-genomes of wheat is unsuprisingly low (3.50%)-methylation levels at these positions are conserved between the genomes. Conversely, at uni-genome methylation sites, it is more common to see a homoeologous SNP between the sub-genomes of wheat that differentiates the methylated genome from the other two sub-genomes (at 65.1% of uni-genome methylated sites). This SNP typically infers a CpG site from a non-CpG site (96% of the time).

### Ancestral methylation can be hard-coded as SNPs

We generated genotype and methylation information for the sub-genome D ancestor (*Ae. tauschii*) to allow comparison with Watkins accessions. We observe that ancestral methylation significantly increases the chance of encountering a different allele in hexaploid bread wheat by ~4-fold (t=-30.42, df=103, p-value<2.2e-16). It shows a predominance for C-to-T/G-to-A transitions that is also statistically significant (t=-283.7129, df=103, p-value<2.2e-16) (Supplementary note 7a, Figure 3b and 3c). These C-to-T/G-to-A transitions are characteristic of the deamination of a methylated cytosine. This apparent preferential deamination of 5-methylcytosine to thymine, has been observed in other organisms^21^ and in *Arabidopsis* where is contributed to bias in spontaneous nucleotide mutation^22^. Furthermore, there was high methylation stability in wheat where most methylation was conserved between *Ae. tauschii* and sub-genome D (83.7%). There was a low level of methylation gain in subgenome D compared to *Ae. tauschii* (3.1%) (Supplementary note 7b).

### Differentially methylated region (DMR) profiles reflect SMP profiles

Gene expression changes are often associated with methylated regions rather than single methylated nucleotides. Using non-overlapping 100bp windows across the genome, DMRs were identified in the CpG, CHG and CHH contexts between each sample and Chinese Spring (Methods)^23^. Per sample, on average 58.7 CpG (range 37-89), 13.4 CHG (range 8-23) and 20.1 CHH DMRs (range 0-168) were identified (Supplementary Table 16). In total 2,356 DMR regions of 100bp were identified across the samples compared to Chinese Spring (491 CpG, 96 CHG and 1,769 CHH DMRs). 1,901 of these DMRs associated with 1,744 genes and 71 DMRs were located in promoter regions associated with 64 genes. For all 2,356 DMR sites, similarly to the analysis for SMP sites, the percentage difference in methylation per sample compared to Chinese Spring was used to cluster the accessions (Supplementary Figure 7). A strong positive correlation exists between the clustering of CpG SMPs and DMRs and as such similar trends are observed with DMRs as was seen for SMPs (Supplementary note 8).

For all accessions, we summarized the number of differentially methylated genes (DMGs) by methylation context i.e. genes with a DMR compared to our reference Chinese Spring (Supplementary Figure 8a). Variation between accessions was highest for CHH DMGs, while the number of genes showing differential methylation in the CpG and CHG contexts is more stable across accessions. CHH variability may reflect the reported dynamic nature of CHH methylation during plant development^24^. There was no evidence of bias in the methylation contexts CpG/CHG/CHH between the wheat A, B and D sub-genomes (supplementary figure 8b-d, and 9a-c).

### Samples cluster by preferentially targeted genes and gene families (Supplementary note 9)

Accessions were clustered based on similarities in the proportion of the number of genes that are methylated in each gene family (vertical dendrogram, Figure 3d). We observe inter-accession variation in gene families highly targeted for methylation. However, a number of gene families are preferentially targeted for methylation across multiple accessions with a high proportion of genes in the family methylated (horizontal dendrogram Figure 3d, coloured red in heatmap). GO enrichment analysis revealed the most common molecular functions associated with highly methylated gene families within and between accessions (Supplementary Tables 17 and 18). Hexokinase activity and glucose binding were the top enriched molecular functions for highly methylated gene families conserved between samples (Supplementary Table 18). These terms are linked to cellular glucose homeostasis and support the hypothesis that some gene families are consistently targeted by methylation across the Watkins collection.

We performed GO enrichment analyses on gene families that were less targeted by methylation within and between accessions (Supplementary Table 19 and 20). NAD binding and N-methyltransferase activity were the top enriched molecular functions for low-level methylated gene families conserved between samples (Supplementary Table 20). Enriched GO terms for highly methylated gene families and less methylated families did not overlap, suggesting that genes of the same molecular function are either consistently methylated or non-methylated across the accessions.

Finally, we focused on CpG methylated genes, appearing in a high, medium or low number of accessions (Methods). Supplementary Figure 10 shows the distribution of the number of genes in the 3 groups: high, medium and low; ~35% of the 2145 CpG methylated genes were present in ≥90 accessions. Previously, we observed that few (0.5%) SMPs were methylated in at least 90% of the samples but this analysis considered CpG and non-CpG sites. For CpG sites, 13.9% of SMPs were methylated in ≥90 samples. Therefore, at the gene level, we see a ~2.5-fold increase in methylation conservation across accessions compared to SMPs (13.9% to 35%). This demonstrates an increased tendency for methylation targeting the same genes across accessions even if the specific cytosine sites differ. Furthermore, the enriched molecular functions within the high, medium and low groups were different with no overlaps (Supplementary Table 21).

### Differential methylation correlates with changes in gene expression

To test the correlation between methylation and gene expression across the Watkins collection, we performed RNA-seq analysis, using 14-day-old wheat seedlings on 12 samples in triplicate, which represent phenotypic tails for; height, heading date, thousand-grain weight and grain width (Supplementary Table 22). We generated gene expression level estimates to allow pairwise comparisons and identify differential gene expression between the samples (Methods). 105,425 wheat genes were analyzed across the sample-set and comparing the 12 samples, 16,112 were differentially expressed; 32.3% from the A-genome, 44.6% from the B-genome and 23.1% from the D-genome (15.3% of analyzed regions) (p-value <0.05).

We normalized allelic gene expression so that per site cumulative expression values for the A, B and D sub-genomes were equal to 100%. The average expression level of sub-genome A across the 289 tri-genome sites associated with promoter regions was 34.22%, sub-genome B 33.43% and sub-genome D 32.35%-demonstrating approximately balanced allelic expression in the sub-genomes. The average expression level of the methylated genome across the 128-promoter associated uni-genome methylation sites was 28.82% while that of the other genomes was on average 35.59%. Therefore, there was a decreased expression of the promoter-methylated sub-genome in comparison to the other two sub-genomes (p<0.0001, t=5.95, df=254).

Previously, we identified DMRs across the samples by comparing non-overlapping 100bp windows with Chinese Spring (Methods). Here, we focused on the 12 samples that were analyzed by RNA-seq and implemented pairwise comparisons to identify DMRs to allow correlation with differential gene expression from the same pairwise comparison. Inter-sample pairwise comparisons yielded an average of; 58.9 CpG, 11.2 CHG and 30.0 CHH DMRs per comparison (Supplementary Table 23). 32.3% of the DMR’s were associated with differentially expressed genes. This reflects a more than 2-fold enrichment in the proportion of genes overall that show differential gene expression. All differentially expressed genes that correlated with DMRs were subjected to the enrichment of molecular functions and biological processes using topGO (p < 0.05) (Supplementary Tables 24, 25 and 26). DMRs that correlate with differential gene expression are more likely to be influencing this expression change and here, CpG DMRs show enrichment for biological processes related to homoeostasis and essential housekeeping. Conversely, non-CpG methylation associates with differentially expressed genes in biological processes related to stimuli response.

For genes that were both differentially expressed and methylated, there is also a bias for enriched GO terms with molecular functions relating to metal ion transportation (Supplementary Table 24). Enrichment for transporter and metal ion binding activity and was seen across SMP sample clusters (Supplementary Table 10 and 11, Figure 1d). This bias of methylation to affect gene expression in pathways related to detoxification and metal ion transportation could be an adaptive response to differences in the soil composition in the country of origin of the sample (Supplementary Table 25, Supplementary note 10). Furthermore, the methylation and gene expression correlations fit the directionality models predicted by previous studies for methylation based on genic positon^25,26,27^. We focused on genes showing differential expression and methylation that had a clearly defined metal ion interaction. This narrowed our analysis to; firstly, a Sodium/hydrogen exchanger that showed up-regulated expression from a (former) Yugoslavian accession 1190352 compared to the Cypriot accession 1190292. Up-regulation of this exchanger is associated with adaptation to salt tolerance that is biologically relevant since Yugoslavia reportedly had large areas of salt-affected soils when Cyprus was at the time unaffected^28,29^. Furthermore, leaves from the Yugoslavian accession 1190352 show significantly higher Na concentrations (average 2182.1 ppm) compared to accession 1190292 (average 1257.7 ppm) (t=5.013, df=4, p-value=0.0074, Supplementary Figure 11a, Methods). Secondly, the ATP-dependent zinc metalloprotease *FTSH 2* showed up-regulation in the Palestinian accession 1190398 compared to a number of other accessions. *FTSH* is down-regulated after exposure of plants to elevated zinc concentrations^30^. Here, the Palestinian accession 1190398 shows *FTSH 2* up-regulation coupled with a lower average leaf Zn concentration (48.63 ppm) compared to each of the three accessions 1190141-China (66.64 ppm), 1190292-Cyprus (68.55 ppm) and 1190352-Yugoslavia (75.28 ppm) for which leaf Zn concentrations were available. The differences in zinc concentrations were not significant however; they fit the directional model for zinc response (t=1.105, df=10, p=0.2949, Supplementary Figure 11b, Methods).

### Early heading date associates with SMP but not SNP profiles

The average expression levels per sample (across the replicates) for the 16,112 differentially expressed genes in one or more of the pairwise sample comparisons, were used for hierarchical clustering (Figure 3e). The barcodes in Figure 3e allow comparison of gene expression clusters with SNP/SMP clusters from Figure 1b and 1d, respectively. Samples that cluster into the same clades by gene expression profiles also cluster closely by SNP profile. This is demonstrated in Figure 3e by conserved colour blocks in the SNP barcode within dendrogram clades. Conversely, samples with divergent expression profiles typically belong to different SNP dendrogram clades. These patterns are also apparent from correlating gene expression and SMP profiles.

Heading date associates with a distinct clustering of samples (Figure 3e). The two samples 1190209/1190034, with earlier heading dates, show the most similar gene expression profiles of all analyzed samples. The samples 1190481/1190181, with later heading dates, cluster together almost as closely but importantly, they are segregated from 1190209/1190034. The two samples with earlier heading dates cluster into the same SMP clade but different SNP clades while, conversely, the samples with later heading dates cluster into the same SNP clade but different SMP clades. This could indicate a common role for methylation in the establishment of an early heading date that correlates with gene expression profile.

We identified differentially expressed genes between early and late heading samples in a pairwise comparison matrix if they were conserved across all replicates; 46 annotated genes were identified (Supplementary Table 27). This includes genes previously linked to flowering time or heading date regulation e.g. REVEILLE 8-like/LHY-CCA1-like 5 that is here down-regulated in early heading date plants^31^. Where methylation associates with these genes, it correlates with the expected directional effect (Supplementary note 11a). Furthermore, Supplementary Table 28 shows the most significantly enriched GO terms and associated biological processes respectively for the 46 differentially expressed genes (topGO, p < 0.05). Enriched processes are predominantly related to meristem growth, development and cell cycle process and phase transition and therefore show biological relevance to the phenotype (Supplementary note 11b).

### Transposable element (TE) abundance is highly variable across the Watkins collection

Analysis of Chinese Spring off-target sequence data demonstrates that it is unbiased sampling of the genome, equivalent to low coverage shotgun sequencing of total wheat DNA, since proportions of TE types closely match those seen in previous shotgun sequence data (Supplementary Table 29, Supplementary note 12a)^32^. To assess TE methylation levels for each Watkins accession, off-target sequencing data was aligned to the wheat TREP database of repeat sequences^33^. Across the Watkins collection, transposons are highly methylated compared to the enriched gene-rich regions (Supplementary note 12b, Supplementary Table 30). This hyper-methylation of repeats is consistent with other plant species and is associated with reducing transposon mobilization.

We observed high variability across the Watkins collection in the proportions of reads aligning to each TE compared to Chinese Spring (Methods, Supplementary note 12c, Figure 4); expansion of retrotransposons is most frequent with 44.2% of accessions showing an increase in mapped base-space of 2% or more compared to Chinese Spring although, large expansions of the mapped base space of 8-10% are seen in DNA transposons in a small subset of lines (Figure 4a). TE expansions do not correlate closely with gene-associated SNP/SMP clusters or geographical clustering. It appears that expansion within the TIR; CACTA group are responsible for increasing the proportion of DNA transposons compared to Chinese Spring in a subset of Watkins accessions (Figure 4b). This expanded group of DNA transposons showed conservation of the high methylation levels seen typically across TEs (Figure 4i). SINE and LTR; Gypsy retrotransposons show prominent and variable expansion compared to Chinese Spring across the Watkins collection (Figure 4c) coupled with conservation of the high methylation levels seen typically across TEs (Figure 4g and 4h). These findings are consistent with previous observations that LTR retrotransposons are epigenetically controlled and a major contributor to genome size change in plants^34^.

**Figure 4.**
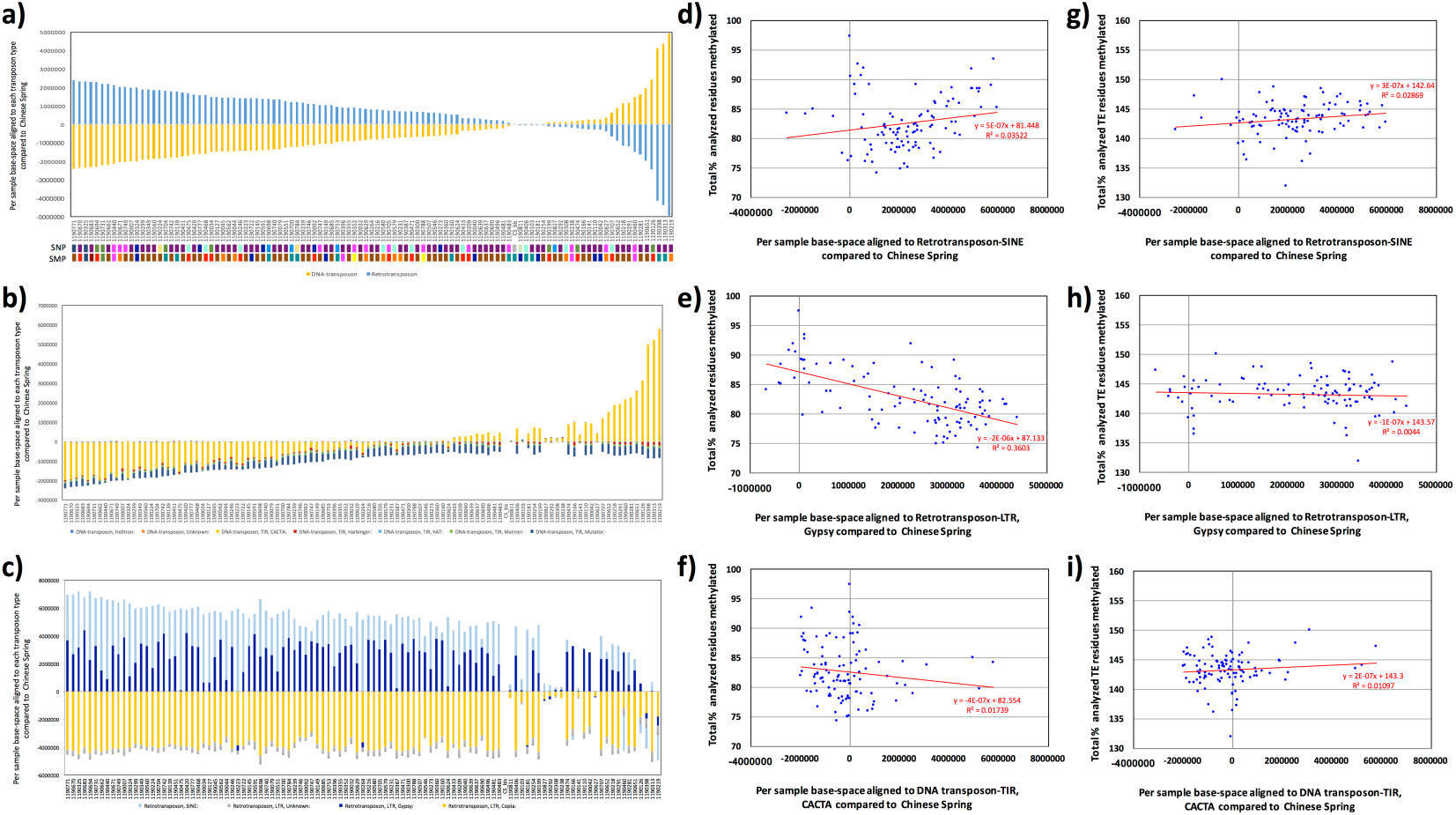
Analyzing transposable element methylation profiles across the Watkins collection. (**a**)Base-space per Watkins accession aligned to DNA-transposons and retrotransposons in comparison to Chinese Spring (Methods) (**b**) Base-space per Watkins accession aligned to DNA-transposons in comparison to Chinese Spring. (**c**) Base-space per Watkins accession aligned to retrotransposons in comparison to Chinese Spring. (**d**) Base-space per Watkins accession aligned to retrotransposon-SINE in comparison to Chinese Spring plotted versus the total cumulative percentages of enriched cytosine residues (gene-associated) that were methylated for CpG, CHG and CHH methylation. (**e**) as per (**d**) but for retrotransposon-LTR, Gypsy (**f**) as per (**d**) but for DNA-transposon-TIR, CACTA (**g**) Base-space per Watkins accession aligned to retrotransposon-SINE in comparison to Chinese Spring plotted versus the total cumulative percentages of TE-associated cytosine residues that were methylated for CpG, CHG and CHH methylation. (**h**) as per (**g**) but for retrotransposon-LTR, Gypsy (**i**) as per (**g**) but for DNA-transposon-TIR, CACTA.

## Conclusions

Using sodium bisulfite treatment and targeted gene enrichment, we observe high epigenomic diversity in the Watkins collection; we demonstrate that methylation is a standalone source of variation in the absence of genetic variation, however, if two wheat accessions show more closely related genotypes then their methylomes are more likely to be related. Both methylation and genotype are influenced by the geographical origin of the sample, although genotypic profiles cluster across wider geographic regions while the methylation profiles of accessions tend to cluster into more local groups. Therefore, we hypothesize that methylation acts as a fast-adaptive response to environmental stimulus. Furthermore, we show that ancestral methylation increases the chance of C-to-T or G-to-A transitions in Chinese Spring wheat that are characteristic of the deamination of a methylated cytosine and may demonstrate this transfer of methylation to SNPs^21,22^. This phenomena could be an important driver of evolutionary change.

We show that tri-genome methylation is more conserved between accessions and therefore the most stable form of methylation, while genome specific methylation sites show enrichment for homoeologous SNPs that differentiate the genome that is methylated from the other sub-genomes. This SNP typically infers a CpG site from a non-CpG site. Tri-genome methylation, correlates with equal expression levels across the 3 sub-genomes while uni-genome methylation correlated with a significant reduction in expression of the affected sub-genome compared to the other two sub-genomes in promoter regions.

Watkins accessions were clustered according to methylation profiles and the clusters show unique profiles of enriched gene function, these variations could contribute to the underlying phenotypic differences between the accessions. Using gene expression analyses, we saw conserved methylation and gene expression profiles in accessions with an early heading date, suggesting that methylation may play a role in the co-ordination of heading date in wheat. DMRs linked directly to gene expression show a bias for genes related to metal ion transportation that links to phenotypic change and could be part of an adaptive response that has been maintained in certain accessions due to differences in the soil composition in the country of origin of the sample.

In addition to epigenomic diversity across the Watkins collection, using Chinese Spring as a baseline, we observe the potential expansion of retrotransposons SINE and LTR; Gypsy most frequently, although some of the largest expansions are seen in a small subset of lines in DNA transposons. These expanded groups of TEs showed conservation of the high methylation levels seen across TEs.

We explore genome-wide epigenetic, alongside genotypic and TE variation across a diverse landrace cultivar collection and open up a new level of genetic variation, which can be exploited by breeders. This provides further opportunities to address important biological questions such as the interaction between epi-type and genotype, the role of epigenetics in the domestication of crops and the stability of and long-term function of methylation in a polyploid genome.

## Acknowledgements

We thank Simon Orford who provided the Watkins seed (BBSRC funded ISP WISP). DNA sequence was generated by The University of Liverpool Centre for Genomic Research (United Kingdom). The enrichment and Illumina sequencing library preparation was performed by LO with support from JK. SNP calling was performed by RJ. The RNA-SEQ work was performed by RRP. JO performed the gene family and gene-ontology analyses with support from MS and KM. The methylation analysis, genotype analysis, manuscript preparation, plant growth and DNA/RNA extraction was performed by LG. The project was designed, planned and conducted by LG and AH. The paper was written by LG and AH with assistance from NH and MB. All authors read and approved the final manuscript. We thank Anita Lucaci and Charlotte Nelson for their assistance with sequencing and library preparation respectively. Sadly, John Danku who performed the ICP-MS analysis for this study died before the data was submitted for publication

## Ethics Approval

Ethics approval was not needed for this study

## Availability of supporting data

All sequencing datasets plus are available (study PRJEB23320) from the European Nucleotide archive (https://www.ebi.ac.uk/ena/submit/sra/#home). Our 12Mb capture design is also available on request.

